# On technical biases in imaging spatial transcriptomics

**DOI:** 10.64898/2026.10.16.759529

**Authors:** Carissa Chen, Lars E. Borm, Anna Pascual-Reguant, Irene Ruano, Davide Risso, Helena L. Crowell

**Affiliations:** Department of Statistical Sciences, University of Padova, Padua, Italy; Center for AI & Computational Biology, VIB, Leuven, Belgium; Center for Neuroscience, VIB-KU Leuven, Leuven, Belgium; Department of Human Genetics, KU Leuven, Leuven, Belgium; Centro Nacional de Análisis Genómico (CNAG), Barcelona, Spain; Universitat de Barcelona (UB), Barcelona, Spain; Padua Center for Network Medicine, University of Padova, Padua, Italy

## Abstract

Imaging spatial transcriptomics (iST) has revolutionized our ability to contextualize gene expression within intact tissue architectures at single-cell and subcellular resolution. However, these platforms introduce complex technical biases that span the entire measurement pipeline – from sample preparation to data acquisition to quantification. Here, we categorize these biases into experimental (e.g., sample preservation and spatial sampling) and computational sources (e.g., z-stack registration and field-of-view stitching). Using public datasets, we demonstrate how different biases can compound and manifest, and propose a unified count modeling framework to formalize these sources of variation. Finally, we call for a more integrated understanding of iST data origins and greater knowledge sharing within and across sectors to support the prospective development of more robust experimental design and reliable downstream analysis.

## Introduction

Spatial transcriptomics enables molecular profiling of tissues *in situ*. Imaging-based approaches (iST) achieve this by microscopic resolution of transcripts in tissue sections through iterative rounds of probe hybridization, imaging, and signal decoding. While some sequencing-based approaches (sST) such as 10x Genomics’ Visium HD and Stereo-Seq [1] have subcellular-sized capture spots, only iST enables reliable single-molecule transcript localization. MERSCOPE (Vizgen; based on [2]), CosMx (Bruker; [3, 4]), and Xenium/Atera (10x Genomics) represent commercial implementations of highly multiplexed 2D RNA imaging workflows, whereas Pyxa (Stellaromics; based on [5, 6]) extends to thicker tissue blocks. All platforms combine platform-specific experimental designs with dedicated image-processing and transcript-quantification pipelines.

Several technical comparison and benchmarking studies have evaluated ST technologies. Technical comparisons primarily describe differences in platform characteristics, such as spatial resolution, molecular coverage, tissue compatibility, and throughput, highlighting practical trade-offs between technologies [7, 8, 9]. Benchmarks extend these comparisons by assessing data quality using standardized samples and evaluation metrics such as transcript recovery, biological accuracy, and reproducibility [10, 11]. These studies provide important guidance for selecting technologies for specific biological applications and motivate further methodological development. Nevertheless, descriptive performance metrics do not fully reveal the variability introduced throughout the iST measurement pipeline. This pipeline involves multiple stages of sample preparation, data acquisition, and preprocessing including tissue harvesting, sectioning, fixation, (optional) tissue clearing, probe design and hybridization, fluorescent imaging, signal decoding, tile stitching, (optional but typical) segmentation, and transcript allocation. Each of these steps has the potential to introduce technical variation influencing downstream analysis and biological interpretation.

While some challenges are specific to imaging-based approaches, they reflect broader statistical and computational difficulties shared across ST technologies. For example, current normalization approaches are problematic for targeted panels in particular [12, 13]; images capture a tissue cross-section from a dynamic 3D architecture; faulty cell segmentation and transcript allocation causes signal ‘bleeding’ between proximal cells [14, 15]; and, imaging-based transcript calling introduces technical noise distinct from sequencing. Failure to call transcripts, for instance, causes false negatives (cf. dropouts in scRNA-seq), while false positives can arise from autofluorescence, probe misbinding, and incomplete stripping of probes. These effects are compounded in higher plex panels that require more cycles of fluorescent staining, probe stripping, and intermittent washes. Increased imaging and overall experimental time can introduce variability in detection efficiency across imaging cycles, can elevate the chance of false positives and negatives, and can negatively affect sample integrity. Together, these sources of variation highlight the need to better catalog and understand technical biases in iST.

Here, we survey technical biases that may arise at each stage of iST data generation (Fig. 1), including histopathology, microscopy, and analysis, in a manner accessible to both experimental and computational researchers. We demonstrate that some biases can be observed in public datasets (Fig. 2), whereas others are either challenging or impossible to validate. In conclusion, we propose that greater exchange of expertise and information between wet and dry labs, as well as across academia and biotechnology, will facilitate the development of more robust experimental protocols and computational tools.

**Figure 1:**
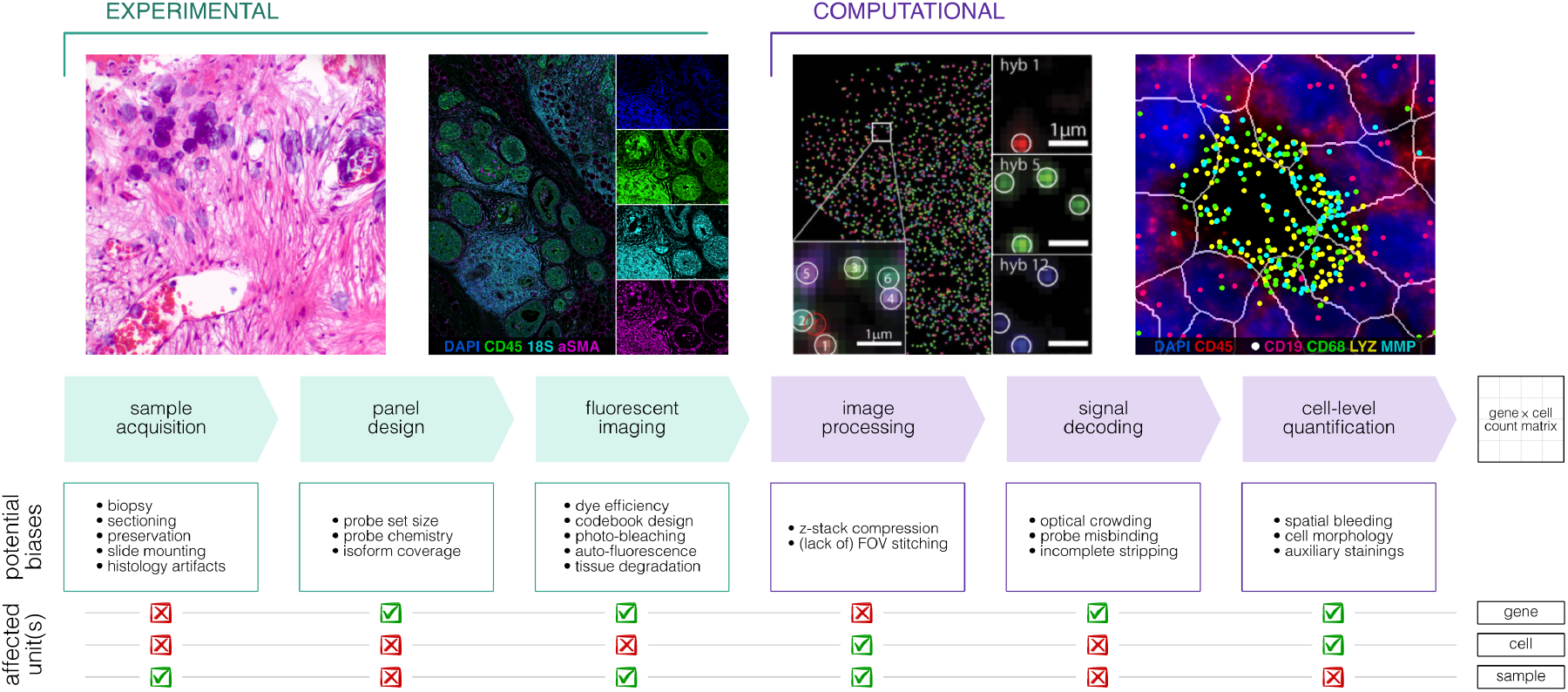
Schematic summarizing experimental and computational steps and biases from sample acquisition to count matrix in imaging spatial transcriptomics. Transforming images into a quantitative gene*×*cell count matrix requires multiple experimental (mint) and computational (lavender) steps, each associated with potential sources of bias that may propagate through the workflow, influence the resulting cellular gene expression profiles, and impact downstream analyses. *[Images modified or produced from mdekmezian, CC0, via Wikimedia Commons; Xenium data on human breast cancer (10x Genomics); Chen et al. [2]; CosMx data on human tonsil [16].]*

**Figure 2:**
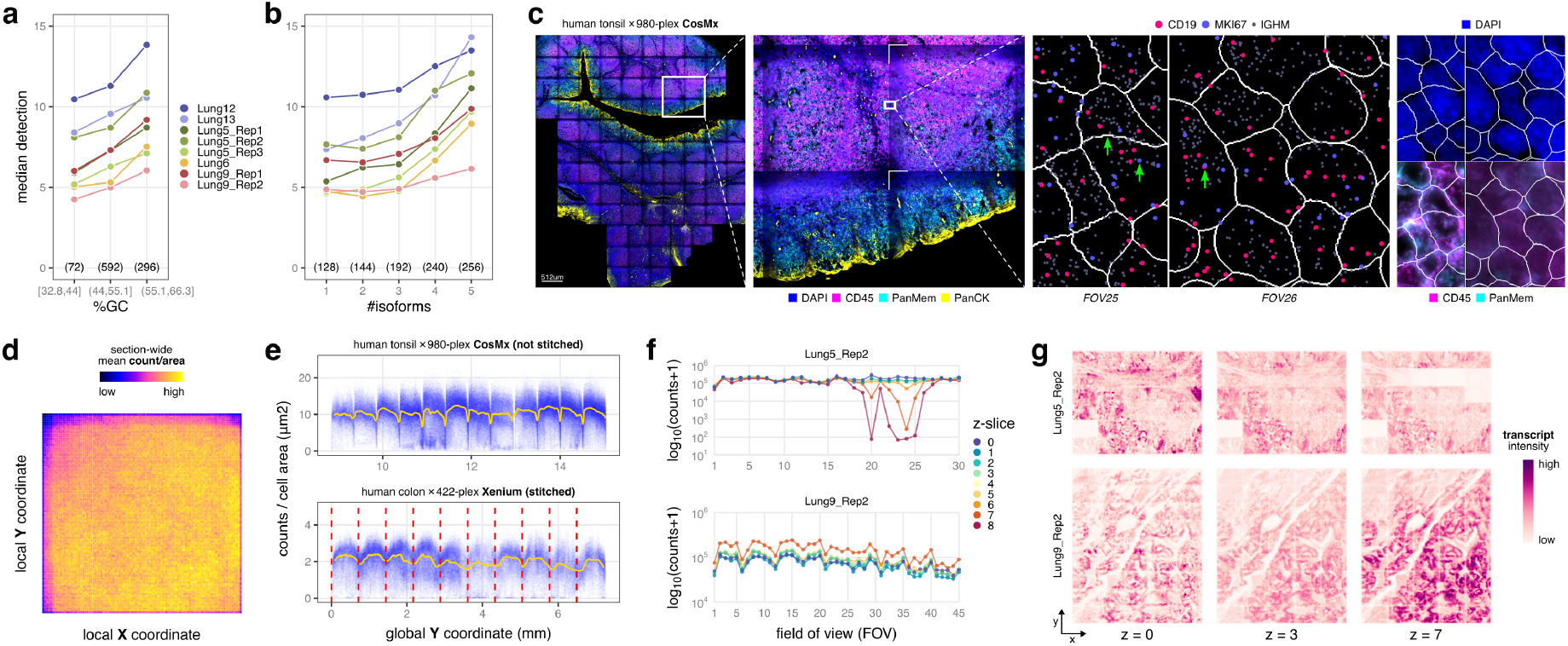
Exemplary technical biases across publicly available datasets. Median percentage of detected targets across cells, stratified by section and **(a)** equal-width bins of mean GC percentage by probe set and **(b)** number of distinct isoforms covered by a probe set; number of targets per x-axis value are shown in parentheses. **(c)** Immunofluorescence composite image including DAPI (nuclei), CD45 (immune), PanMem(brane) and PanCK (epithelia); zoom-in on four fields of view (FOVs); and, zoom-in on neighboring FOVs with CD19, MKI67, and IGHM transcript molecules, highlighting discrepancies in cell segmentation and transcript allocation. **(d)** Heatmap of RNA density averaged within 100 *×* 100 local coordinate bins across all FOVs in the tissue section. **(e)** RNA transcript counts per area (px) vs. distance (mm) to field of view (FOV) borders; blue points = cells; yellow line = rolling mean (window size 20). **(f)** RNA target counts stratified by FOV and z-slice for two exemplary tissue sections. **(g)** Corresponding transcript densities visualized spatially; rows = sections, columns = z-slices. CosMx data are from [16] and [3], Xenium data from [51].

## Experimental: from samples to images

### Sample preservation, sectioning & slide preparation

Tissues for iST are typically fresh-frozen (FF) or formalin-fixed paraffin-embedded (FFPE). Preservation by FF offers superior RNA quality but is more prone to the formation of ice crystals that disrupt cellular morphology [17, 18]. FFPE, the clinical standard with decades of archival material, provides excellent morphological preservation but at a cost of RNA quality due to room-temperature storage and overfixation, which can reduce hybridization efficiency [17, 19, 20]. Independently, preanalytical variation can arise before sectioning, during biopsy collection, fixation, and embedding. Cold ischemia time, fixation delay and duration, tissue processing conditions, and prolonged storage of tissue blocks (or pre-cut sections) can affect RNA integrity and accessibility [21]. RNA quality may have platform-dependent effects downstream: a recent study found that lower DV200 values might be associated with reduced transcript diversity and counts in Xenium, whereas no detectable association was observed for CosMx [22].

During sectioning, common histological artifacts include blade chatter and compression, tissue folds and wrinkles, physical tears and detachment, and stress-induced distortions [23]. Furthermore, sectioning – typically at 5-10 *µ*m – slices through cellular volumes, yielding a 2D profile in which portions of the cell (and their transcripts) are discarded. Section thickness therefore influences what fraction of a cell is captured and, consequently, the expected transcript count and degree of (vertical) contamination per cell [11].

After sample preparation, iST vendors provide standardized, one-size-fits-all slide preparation protocols. However, critical steps can strongly influence data quality and vary substantially between tissue types. Experimental conditions during, for example, tissue decrosslinking and permeabilization, fiducial incubation, photobleaching, and probe hybridization are often left to users, with few studies optimizing and reporting them. Because iST relies directly on microscopic visualization of fluorescent readouts, these artifacts have the potential to propagate into downstream analyses. An understanding of the prevalence, characteristics, and potential impact of such artifacts is therefore also relevant for bioinformaticians.

### FOV placement, replication & spatial sampling

The portion of a sample that can be imaged in an iST experiment is limited by a pre-defined sample area, varying between several square millimeters to centimeters depending on vendor, where the user has to place their tissue. In addition, the user can select regions of interest (ROIs). To resolve individual RNA molecules, the microscopes used for iST require sufficient magnification and resolution, which comes at the cost of small fields of view (FOVs), so that the ROI has to be further subdivided into many FOVs, typically arranged in a grid. ROI and FOV placement may be guided by the biological question at hand as well as histology (e.g., avoiding necrosis, folds, tears, or tissue loss). Imaging all FOVs for each color channel, z-plane, and cycle (determined by gene panel plexity) takes the bulk of acquisition time. Prolonged imaging could harm sample integrity, likely due to the generation of photo-induced reactive oxygen species (ROS) [24]. The finite capture area per slide and per-slide acquisition time thus cap spatial coverage per sample and overall sample throughput, respectively.

A prominent experimental design strategy is provided by tissue microarrays (TMAs), which enable comparative spatial studies [25, 26, 27]. By arranging dozens of tissue cores on a single slide, TMAs increase cohort size while reducing technical variability. To support quality control and normalization, TMA experiments commonly include reference tissues or cell line controls, which can also serve as physical orientation guides across slides and batches. However, they inherently amplify spatial sampling bias: cores are typically 0.6-3 mm in diameter, which may be insufficient to fully represent tissue heterogeneity and can miss rare or spatially localized anatomical structures and subpopulations [28]. Besides, individual cores are extracted using a hollow punch tip, resulting in mechanical disruption of the tissue at the core margins. The cores used in TMAs, as well as small tissue biopsies, are further prone to detachment during slide preparation and data acquisition due to their limited surface contact and support from surrounding embedding material. Core diameter and spacing also impose practical constraints, as they must be compatible with the capture area limitations of downstream iST platforms.

Spatial sampling arguably manifests across several scales: the profiled individual, biopsy site, tissue section, and its measured portion. This area should cover the spatial variation present in the sample, which can be challenging for large specimens (e.g., human tissues), or when only small pieces can be sampled (e.g., biopsies). Larger sections or measurement areas can mitigate spatial sampling biases, but at the cost of biological replicates and longer experiments. Vice versa, TMAs ease replication but amplify spatial sampling biases. These effects are particularly relevant when comparing cell types, histological regions, or biological samples: differences in composition or morphology can stem from sampling variability, independent of true molecular variation. While experimental design has received limited attention in ST [29], distinguishing genuine biological organization and differences from artifacts requires careful consideration of both sample selection and measurement processes. Thus, iST experiments require a trade-off between sample throughput, representativeness, and data quality, while careful study design, perhaps supported by simulations, remains critical [30, 31].

### Probe sequences, sets & panel design

Commercial iST platforms include three types of barcodes, corresponding to either a target probe, a negative control, or neither (Box 1). Probe sequences introduce four potential sources of bias: length, GC content, self-hybridization, and (non-)complementarity to off-target sequences. To mitigate such biases, iST platforms employ probe-design strategies that constrain probe characteristics while avoiding strong predicted secondary structures and regions of high sequence similarity. In sST, homologue spot sequences were found to produce systematically lower-quality observations across slides and tissue types [32]; off-target binding has likewise been observed in iST [33]. Biases associated with fragment length and GC content has been highlighted in RNA-seq [34] and sST [35], while probe length appears to have limited effects in iST [36].

However, probe sequences and sets (i.e., the different probes used to target each gene) are mostly proprietary, particularly for pre-designed panels that dominate current applications. An exception is the company preprint accompanying the pre-commercial 1K CosMx human NSCLC dataset [3], which provides both probe sequences and fluorescent barcodes. The majority of probes are 35 bps in length, and GC content is approximately normally distributed across probes (mean and median ∼52%). Consistent with sequence-dependent differences in hybridization efficiency, genes targeted by probe sets with higher GC content show increased detection (Fig. 2a). Lack of publicly available information, however, complicates systematic evaluation of probe- and target-specific biases associated with assay design across platforms, panels, and tissues.

### Alternative splicing & isoform selection

While iST reports transcript detection at the gene level, it relies on predefined probe sets that target specific untranslated regions, exons or exon-exon junctions. Targeting short transcript regions with gene-specific oligonucleotide probes allows for detection of RNA despite fragmentation and, importantly, provides retrospective access to FFPE sample archives. Alternative splicing, however, is a ubiquitous biological mechanism that produces multiple transcript isoforms from a single gene. If a tissue or specific cell type expresses an isoform that lacks the targeted exons, it will be systematically under-detected or reported as absent (false negative). Moreover, probes spanning an exon-exon junction that is spliced out in certain cellular states will fail to bind. This mismatch between panel design and biological isoform variety may introduce cell type-specific quantification biases that are independent of relative gene expression levels. Of note, methods to infer isoform usage from ST data [37] and to spatially profile transcript isoforms [38] have only recently been proposed, while isoform-level resolution is also limited in widely studied single-cell and -nucleus RNA-seq data based on short-read or probe-based sequencing assays [39].

In the He *et al*. [3] panel (960 targets, five probes each), 828 genes (86.3%) have differential probe coverage across annotated isoforms. In fact, only 502 genes (52.3%) have all their isoforms covered by at least one probe, while only 100 genes have every isoform covered by all five probes (Ensembl hg38 annotation v116). Collectively, the panel targets genes with 22,834 annotated transcripts, of which only 12,767 are fully covered by all five probes and 2,055 have no exact probe matching. Furthermore, the median detection increases with the number of distinct Ensembl isoform sets covered by the probes (Fig. 2b). Although this pattern is consistent with broader isoform coverage improving gene detection, it does not establish a causal relationship. In particular, the association may be confounded by the number of annotated isoforms per gene, because genes with more annotated isoforms can generate more distinct probe-coverage patterns and may differ systematically in their expression or degree of annotation.

We note that for 927 genes (97.5%), the MANE Select transcript [40] is matched by all five probes. This indicates that incomplete coverage is disproportionately concentrated among alternative and minor transcripts. However, recent evidence from long-read sequencing data suggests that alternative splicing is tissue- and disease-specific [41, 42]. Approximately half of all expressed genes in GTEx display tissue-dependent transcript isoform expression [43], and thousands of transcripts are exclusively expressed in a single tissue or have different transcript ratios across tissues [44]. Hence, when isoform usage differs between cell types or conditions, iST can report different gene-level expression even when the total number of gene molecules is unchanged. Since commercial platforms do not publish probe sequences, however, our analysis is limited to a single pre-commercial dataset [3].

### Fluorescent barcoding, optical crowding & autofluorescence

Detection of RNA in iST relies on error-robust, barcoded fluorescent *in situ* hybridization: each target is assigned a temporal barcode that is decoded from the pattern of fluorophore signals observed across imaging cycles [2]. As mentioned above, the encoding scheme typically includes three classes of barcodes: target probes are complementary to an RNA sequence, non-complementary negative or control probes reflect untargeted binding, and false or blank codes without associated probes reflect decoding accuracy. Each barcode is defined by the positions and fluorophores of its positive bits and is, by design, sparse and mutually dissimilar, conferring resilience to single-bit errors (Box 1). Because detection efficiency varies with both dye identity and imaging cycles, fluorescent barcodes introduce gene-specific detection biases. Specifically, differences in dye brightness and stability (i.e., resistance to photobleaching) lead to variations in detection sensitivity and false positives. Sample deterioration and photobleaching over imaging rounds exacerbate these biases, as barcodes with positive bits positioned in later cycles are more prone to signal loss.

Transcript density inside the cell is higher than can be resolved by diffraction-limited microscopy, so only a subset of genes should be simultaneously labeled. This optical crowding impairs the accurate resolution of individual fluorescent spots and, thereby, the decoding of fluorescent barcodes when too many genes with highly colocalizing transcripts are visualized in the same barcode bit [45, 46]. Proximal spots, for instance, may be identified as a single signal, leading to local count saturation. This compression of the dynamic range means that differences in high-abundance transcripts are systematically underestimated and, worse, can negatively affect other genes labeled in the same barcode bit. Dense transcript areas can also inflate the false discovery rate (FDR) because crowded signals can appear like off-target barcodes by chance. The trade-off between multiplexing capacity and error robustness was highlighted by Chen *et al*. [2]. Dense encoding schemes [47] maximize the number of resolvable targets but suffer from reduced calling accuracy and higher misidentification rates as panel plexity grows. In contrast, sparse schemes trade off target capacity and barcode length for sparser signal and, thus, improved detection efficiency. The FDR for rare molecules can be reduced by optimizing codebook and decoding algorithm design (e.g., by incorporating priors on transcript abundances [48]).

Beyond multiplexing-related challenges, biological tissues can exhibit autofluorescence with broad emission spectra [49]. Common sources include paraformaldehyde overfixation and several naturally present components (e.g., lipofuscin, heme derivatives, metabolic cofactors such as NAD(P)H, extracellular matrix (ECM) components such as collagen and elastin) [50]. Tissue autofluorescence increases and spatially structures background fluorescence, thereby reducing the signal-to-noise ratio (SNR) and causing channel-specific, inter-sample, and regional SNR differences. Autofluorescence varies with tissue type and condition as well as wavelength, generally affecting channels in the middle of the visual spectrum most strongly, with liver, spleen, lung and skin showing particularly high levels. FFPE tissue often exhibits higher autofluorescence than FF tissue as aldehyde fixation can increase background [11]. And, differences in metabolic or ECM composition contribute to variability in autofluorescence across and within tissues (e.g., lipofuscin may accumulate with age and disease, ECM deposition during inflammation and fibrosis [50]).

Although recent advances have markedly increased plexity, the number of ‘trustworthy’ observations across all targets ultimately saturates due to detection limits, spot-recovery inefficiencies, and decoding constraints. For fixed-length barcodes, the number of reliably resolved spots declines as plexity increases because read-out errors accumulate non-uniformly, with molecular crowding disproportionately affecting highly abundant transcripts and transcriptionally dense regions. Moreover, commercial platforms keep their encoding and error-correction strategies proprietary. So while understanding these biases is valuable, the lack of disclosed information limits their direct evaluation and, consequently, our ability to model and correct for them.

### Wide-field epifluorescence microscopy & temporal scanning sequence

The optics system in a typical iST instrument, such as CosMx, MERSCOPE, and Xenium, uses wide-field epifluorescence microscopy [3, 52] where the FOV is illuminated at once to excite fluorophores at designated wavelengths, and the emitted light is separated into different color channels using optical filters [53, 54]. The microscope objective and associated optical path can cause field-dependent illumination, including non-uniform illumination and vignetting [55], which often causes a radial decline in excitation and fluorescence detection toward the FOV periphery. If not adequately corrected for, transcripts near the FOV edge are subject to systematic under-detection compared to those in the center. In addition, wide-field epifluorescence microscopy can introduce other non-uniformities, including optical aberrations that vary across the FOV and with depth, wavelength-dependent shifts in focus between channels (chromatic aberrations), and background from fluorescence above and below the focal plane [53, 56, 57].

The sequential acquisition of FOVs could further introduce a temporal axis of variation. Over the course of an experiment, reagents such as ROS scavengers deplete, buffer components age, and the sample is exposed to repeated cycles of bleaching and laser excitation. However, the lack of temporal scanning information and, in some cases, FOV positioning information, precludes quantification of these effects in commercial iST data. In all, spot detection probability is not uniform across a slide, but may be influenced by relative FOV position (center/edge), acquisition order (early/late), imaging channel (wavelength), and tissue depth.

## Computational: from images to counts

### FOV stitching, z-stack registration & spot-calling

Depending on their placement, FOVs might (not) overlap and can be stitched into a smooth composite image. Stitching can lead to duplication of molecules when registration is imperfect, whereas lack of stitching can yield fractured and duplicated segmented cells. In CosMx, where cell segmentation is performed per FOV (without stitching) [3], seemingly identical cells can exhibit data discrepancies (Fig. 2c). In addition, a drop in transcript counts can be observed at FOV borders (Fig. 2d). Although Xenium performs stitching prior to cell segmentation, these effects remain observable for both platforms (Fig. 2e).

While many downstream analyses rely on a flattened 2D image, tissue sections are cut at a defined thickness and, therefore, retain an inherently 3D perspective with cells, organelles, and other tissue structures distributed across different z-planes. To provide a more complete representation of individual cells for segmentation, and to capture molecular signals throughout the tissue, imaging entails acquiring a sequence of 2D images along the z-plane at discrete depths, so-called z-stacks. However, z-stack acquisition introduces several technical artifacts. Uneven tissue sectioning, for example, results in an axial tilt. The non-uniform refractive index of tissue could also cause depth-dependent optical effects. As the microscope focuses deeper into the tissue, light attenuation and scattering may occur as photons are absorbed or scattered by other obstructing molecules, reducing overall signal intensity [58]. Extended acquisition time may also lead to increased photobleaching [59]. Collectively, these effects may result in uneven fluorescence quantification throughout the tissue (Fig. 2f,g).

### Defining analytical units & cell segmentation

Although iST offers molecular resolution, the main quantity of interest for downstream analysis is typically a count matrix obtained by performing ‘instance’ segmentation (cf. ‘semantic’ segmentation, which distinguishes cells from background but not one another) and counting gene-wise transcripts in each instance. Gene-by-instance counts thus depend strongly on the data modalities available, the segmentation strategy employed, and what an instance represents (e.g., cell nuclei or bodies, peaks in transcript density).

Available modalities can vary between commercial platforms. In general, immunofluorescence stainings aim to provide complementary information on nuclear and cellular morphology, and cell identity [60], with DAPI used to identify nuclei as segmentation anchors [61]. Depending on the sectioning plane, however, a cell may be intersected without its nucleus, and multi-nucleated, syncytial or adjoining nuclei can complicate one-to-one nucleus-cell correspondence [62]. Nuclei thus provide an imperfect representation of cells, and their segmentation inevitably introduces uncertainty before cell boundaries are estimated. Isotropic, radial, or Voronoi-like expansion can be applied to approximate whole cells, whereas cytoplasmic RNA signals (e.g., poly(A), 18S rRNA) and membrane markers (e.g., ATP1A1, CD298/B2M) can provide more faithful information on cellular extent and boundaries, respectively. Cell type-associated stainings, such as CD3, CD45 and CD68 (immune), *α*SMA and Vimentin (stromal), or PanCK and E-cadherin (epithelial), can further aid segmentation [63, 64].

The choice of staining and segmentation strategy therefore directly influences segmentation accuracy, the fraction of transcripts assigned to each cell, and potential biases. Weak or absent membrane staining can lead to conservative segmentations that exclude a substantial fraction of transcripts. Conversely, heuristic boundary expansion can increase signal contamination and distort cellular morphologies. Accurate segmentation is particularly challenging in tissues lacking reliable membrane markers (e.g., adipose tissue) or exhibiting complex cellular morphologies (e.g., neurons) [65, 66, 67]. Across a variety of datasets, 39% of transcripts have been reported to remain unassigned on average, with contributions from lateral diffusion and technical noise (false positives) besides faulty segmentation (missing cells) [68].

Many transcripts also exhibit compartment-specific localization. Mature mRNAs are predominantly cytoplasmic, whereas nascent transcripts, retained intronic RNA, and many long non-coding RNAs are enriched in the nucleus [69, 70, 71]. Specialized for iST, more recent approaches let transcripts inform segmentation, albeit at the risk of not capturing cellular morphologies faithfully [72, 73, 74, 75]. At the end of the spectrum, cell segmentation-free approaches tackle the segmentation challenge by relying solely on transcripts [76, 77], although their adoption remains limited, perhaps owing to limited interpretability and heavy computational demands for very high-plex data. Overall, nuclear-centered, whole-cell, transcript-informed, and cell-free approaches define different units of analysis with systematically different transcriptomes, thereby affecting all downstream ‘cell’-level analyses.

### Transcript (mis)allocation, decontamination & modality differences

Misallocation of transcripts is a primary driver of technical noise in iST, arising from both computational and biological sources. Optical imaging projects 3D tissue structures onto a 2D coordinate system, either before segmentation (e.g., max-projection of fluorescence intensity) or after segmentation (e.g., projecting segmentation masks). This projection collapses vertically adjacent features, increasing signal overlap and complicating 2D boundary delineation [78]. Intertwined with segmentation, several physical and optical factors contribute to transcript misallocation, or spatial ‘bleeding’, between proximal cells.

First, vertical overlap between partial volumes causes cells lying immediately above or below the sectioning plane to bleed transcripts into the masks of neighboring cells [14]. Second, complex, entangled, or branching cell morphologies (e.g., thin processes of neurons, astrocytes, or myeloid cells) are extremely difficult to capture with standard boundary-finding algorithms; transcripts located in these extensions are frequently misassigned to the cell bodies of nearby neighbors. Finally, transcripts can undergo physical displacement during sample preparation, leading to molecules being called outside their cell of origin. As a result of these errors, single-cell profiles represent a mixture of a cell’s own transcriptome and that of its local microenvironment. This contamination compromises all stages of downstream cell-level analysis, including dimensionality reduction, cell-type annotation, differential expression analysis, and cell-cell communication inference [14, 78].

While computational remedies remain sparse, several have been proposed. For example, *Statial* [79] uses cell-type deconvolution estimates to account for signal contamination during statistical testing. Other approaches try to refine segmentation boundaries and per-cell transcriptomes, or to address transcript misallocation after cell segmentation. These include boundary refinement tools such as *FastReseg* [80] and *segger* [81], cell-simulation approaches like *Proseg* [75], deconvolution-based frameworks like *SPLIT* [82], and transcript-level denoising tools like *ResoIVI* [83] and *DenoIST* [84]. As highlighted by Ishaque *et al*. [78], however, evaluating these tools remains challenging due to a lack of ground-truth data and standardized evaluation metrics. Reliable cell segmentation and transcript allocation thus remain central bottlenecks in iST data analysis. Emerging 3D data modalities (e.g., [5, 85]) could help mitigate spatial bleeding, but introduce new segmentation challenges and require substantial computational resources. And, acquisition of such data remains time-intensive and costly, limiting scalability and routine application.

Some computational tasks (e.g., cell-type annotation [86, 87], contamination modeling [14, 80, 82, 81]) either rely or improve upon integrating iST data with single-cell or -nucleus RNA sequencing (scRNA- and snRNA-seq, respectively) reference data on matched samples or from large-scale atlases [88, 89]. However, these assays introduce a bias in the cell types and transcripts recovered [90, 91]. Unlike iST, scRNA-seq requires enzymatic and mechanical dissociation of intact tissue into a suspension of viable cells. Fragile, highly adherent, morphologically complex, or ECM-associated cells are particularly susceptible to loss, whereas immune and other readily dissociated cells may be overrepresented [92, 93, 94]. And, compared with scRNA-seq, snRNA-seq preferentially detects unspliced transcripts, intronic reads, and nuclear RNAs while under-representing many cytoplasmic transcripts.

Thus, besides biological variation, different cell type abundances and their transcriptomes have well-understood technical origins, independent of how analytical units are defined. Differential transcript localization, delineation of analytical units, and cell-type recovery efficiency will jointly manifest in downstream analyses. Single-cell references, for instance, may not accurately capture iST-derived gene expression; and, underrepresented or missing populations may be annotated with reduced confidence, contribute less to deconvolution, or be overlooked during contamination correction. Any such effects should therefore be considered when interpreting data from iST or when integrating data across modalities.

### Count modeling & statistical inference

The main object of many downstream analyses is a gene-by-cell count matrix. Hence, it is common in the bioinformatics and statistical literature to equate these data to single-cell sequencing assays, and methods from scRNA-seq are often used or adapted by including the spatial location of each cell, typically its centroid. The standard scRNA-seq measurement model consists of a Poisson process in which a fixed number of total reads (a.k.a. *sequencing depth*) are distributed among genes in proportion to their relative expression [95]. When including biological variation in expression, this leads to a negative binomial model with a cell-specific offset, its *library size* [96], justifying the conventional normalization approach of scaling by total count.

RNA-seq measurement models do not have a sound justification in iST assays. Instead, a proper measurement model can be defined by assuming a Poisson point process [97]: defining cells as a finite set of polygons, and assuming constant intensity of the process within each cell, leads to a Poisson distribution. Further assuming that the intensity of the process for each gene is modeled by a Gamma mixture model across cells [75] leads to a negative binomial marginal distribution, with mean given by the product of expression intensity and cell area (Box 2). Taken together, this derivation leads to a negative binomial model similar to that of sequencing data, but where the offset relates to the relative size of each cell rather than the total read or transcript count (see also [12]).

Due to segmentation variations and differences in 3D cell thickness across the slice, segmented cell areas and volumes vary by orders of magnitude. Larger segmentations naturally accumulate more transcript counts, but may reduce transcript density (i.e., counts per area). If normalization is done by simple total-count scaling (adapted from scRNA-seq) or naive division by cell area, mathematical artifacts are introduced. Especially for lower-plex panels, context-tailored gene selection may lead to confounding between total count and biological signal [12]. The variance in cell size remains a massive downstream confounder that risks misidentifying distinct cell states (e.g., active vs. inactive) during clustering.

The mean of the negative binomial distribution represents the expression of each gene in each cell. This mean can be modeled as a function of several parameters, which represent the biological and technical factors that influence gene expression in iST (Box 3). Here, we take a comprehensive view, in which we list all possible gene- and cell-specific effects that contribute to cell-level gene counts, even though not all of them will be identifiable from experimental data due to confounding. Nonetheless, enumerating these effects can help inform the design of new benchmarking datasets and normalization strategies.

Specifically, we decompose the mean expression in additive terms that correspond to i) biological transcript intensity; ii) technical detection efficiency; and, iii) geometric sampling exposure. The biological component is influenced by each cell’s total amount of RNA, each gene transcript’s true relative abundance, the effect of a latent cell class (e.g., cell type or state), and the effect of observed biological covariates (e.g., genotype, cell-cycle phase, disease status). We also include a smooth spatially varying biological effect (such as a phase-transition gradient) as well as a cell’s microenvironment effect, assuming that gene expression is affected by neighboring cells. The technical component includes intra- and inter-FOV variation as well as gene-specific probe effects, such as barcode-decoding sensitivity and hybridization efficiency, including GC-content effects and probe-isoform matching. Finally, as noted by [75], cross-cell contamination due to segmentation errors can be a further source of technical variation (see Box 3 for technical details).

Since the inception of iST, countless methods have been proposed for its analysis [98, 99], ranging from the identification of spatially variable genes [100, 101], to spatial clustering [102, 103], to differential expression [104, 105]. However, several computational and statistical challenges are still outstanding. For instance, many gene prioritization and clustering approaches employ Euclidean distances to model spatial relationships among transcripts and cells. This is problematic as tissue geometry is not isotropic (i.e., the interpretation of distance between cells is not the same in all directions), especially in the presence of blood vessels, complex organ structures, or at the cancer-stroma interface. While some deep learning- and spatial network-based methods go beyond the Euclidean space assumption [106, 107], methods developed for non-Euclidean geometry in other fields may be a promising avenue for future research [108].

As for differential expression, many analytical workflows employ approaches borrowed from scRNA-seq. While methods have been proposed to account for spatial dependencies among cells [109, 110, 111], these are not yet widely used and suffer from the same Euclidean geometry problem discussed above. Furthermore, when comparing multiple samples across patients and disease status, there is a risk of confounding association between gene expression or cell-type composition with the spatial sampling of different tissue specimens: indeed, we typically assay a very small portion of a sample, and what can be directly compared across subjects may be biased by uneven ROI selection. As stated above, careful experimental design – which and how many areas of a sample to assay per individual – is paramount.

Taken together, trustworthy iST analysis needs measurement models, normalization strategies, and spatial statistics built around the geometry of the tissue rather than borrowed from single-cell sequencing, in order to separate genuine biological signal from technical artifacts. Furthermore, true population-level inference requires independent biological replicates, at the individual, rather than cell or FOV, level. This is especially important in populations with high biological variation (e.g., considering humans *versus* genetically identical mice), where a large number of biological replicates with consistent spatial sampling is needed to justify popular statistical measures, such as *p*-values, and to support robust scientific conclusions.

## Discussion

The biases cataloged here span the full iST data generating processing, from experimental acquisition to computational processing, yet they differ in what an analyst can do about them. Some represent moderate biases, such as optical vignetting or registration issues, that could be measured, partially corrected for, or modeled as covariates. Others represent outright loss of information: sectioning artifacts, gross tissue damage, severe de-focus, failed FOVs, etc. These can not be modeled away, but could be flagged or masked during quality assessment.

Several computational tools have been developed for quality control (QC) of different iST data layers, including cell-level count matrices [112], single molecules [113], histology stainings [114], IF images [115], or a combination thereof [116]. Upcoming tools represent more iST-tailored QC frameworks to assess cell segmentation, transcript allocation, or imaging quality (e.g., *SegTraQ* and *spoQC*). However, QC aims to flag, remove, or correct problematic data – be it cells, expression profiles, tissue regions, or whole sections. It does not, nor does it aim to, guarantee that technical biases are accounted for during downstream statistical modeling and inference.

Intrinsic to the wide-field epifluorescence design, there are imaging artifacts that become especially relevant when tiling hundreds of FOVs to cover whole tissue sections. These are increasingly recognized, but are not yet comprehensively defined, standardized, or transparently corrected for across commercial iST platforms. The extent and performance of optical correction (e.g., field-dependent illumination, point spread function variation, chromatic misregistration, or tile-boundary effects) are rarely available to users as assay-level QC metrics or correction parameters.

Technical challenges are amplified in tissues that require specialized processing conditions. For example, bone commonly requires decalcification, which can affect RNA integrity and probe accessibility [117]; brain tissue is highly sensitive to post-mortem changes and to variation in perfusion or fixation conditions, which can increase autofluorescence and affect RNA preservation or tissue morphology [118]; and, adipose tissue is particularly difficult to section and adhere to the slide. Moreover, FFPE preservation dissolves intracellular fat, leaving empty adipocyte vacuoles that complicate segmentation and transcript assignment [65]. Technical constraints such as microscope resolution, cell type-specific IF stains, and limited applicability to non-mouse or non-human specimens (e.g., plants [119] and prokaryotes [120]) further complicate applicability of iST.

The principal experimental and computational steps of the biotechnology underlying, e.g., CosMx (Bruker), MERSCOPE (Vizgen), and Xenium/Atera (10x Genomics) are known. Yet, the level of default metadata reported varies, and many technical details remain undisclosed. Standard CosMx and MERSCOPE outputs include FOV assignments as cell metadata, whereas Xenium provides them for transcripts, with FOV coordinates available in auxiliary files; the order in which FOVs were acquired is generally unknown. Run-, transcript- and cell-level QC metrics vary substantially in their availability, comprehensiveness, and utility across pipelines. Proprietary workflows further withhold critical information including probe sequences, barcode catalogues, reagent concentrations, and imaging parameters. In addition, vendor pipelines typically include image registration, feature detection, cross-round decoding, and quality filtering, among other steps. Taken together, this lack of transparency creates a widening gap between biotechnological development by industry, on the one hand, and computational analysts, on the other. As a result, researchers must devise robust analytical methods without full insight into the data generating process.

Addressing technical biases is essential for translating spatial molecular maps into robust biological insight. Achieving this will require greater transparency, shared standards, and closer collaboration among biotechnology companies, experimentalists, and computational scientists. In particular, curated, metadata-complete iST datasets could provide a foundation for systematically benchmarking bias-correction strategies in a vendor-neutral manner. By systematically surveying technical variation and bias in iST, we aim to foster greater transparency and methodological rigor in the field.

## Code & data availability

Analyses were run in R v4.6.0, with Bioconductor v3.23 [121]. Code to reproduce Fig. 2 is accessible at https://github.com/HelenaLC/ist-biases; analyses underlying panel (b) were supported by Codex with GPT-5.6 Sol (see corresponding commits). The CosMx data on human tonsil are available from Crowell *et al*. [16], CosMx data on human NSCLC from https://brukerspatialbiology.com [3], and Xenium data on human CRC from https://www.10xgenomics.com [51].

## Acknowledgements

L.E.B was funded by the EU’s Horizon Europe research and innovation program under Marie Sklodowska-Curie grant agreement N°101146749. A.P.R. receives funding from the MCIN/AEI/10.13039/501100011033 and FSE+ (RYC2022-035848-I) and the MICIU/AEI/10.13039/501100011033/FEDER/UE (PID2023-148687OB-I00). D.R. was supported by the European Research Council (ERC; Grant CoG 101171662). H.L.C. was funded by the Swiss National Science Foundation (SNSF; project grant number 222136) during part of the project period.

## Authors’ contributions

Author contributions following CRediT taxonomy: *Conceptualization*: DR, HLC; *Formal analysis*: CC, DR, HLC; *Funding acquisition*: APR, DR, HLC; *Supervision*: DR, HLC; *Visualization*: CC, HLC; *Writing – original draft*: DR, HLC; *Writing – review & editing* : CC, APR, LEB, IR.

## Declaration of generative AI and AI-assisted technologies in the writing process

During the preparation of this work, the author(s) used Claude Sonnet 4.6 and ChatGPT-5.6 Luna to assist with proofreading, readability, and concision. After using these tools, the authors reviewed and edited the content as needed and take full responsibility for the content of the published article.

### Box 1: Glossary of imaging spatial transcriptomics

**Target probe:** a short oligonucleotide (10s of nts) complementary to a specific molecular target (e.g., transcript region), containing an encoded fluorescent reporter-binding sequence.

**Target probe set:** a library of transcript-specific probes, designed to hybridize different regions (often different isoforms) of the same RNA target. This redundancy improves detection sensitivity and robustness while reducing the impact of individual probe failures or non-specific interactions.

**Negative probe:** an oligonucleotide probe designed with similar molecular properties as target probes but lacking complementarity to the sample transcriptome. Detection of these probes provides an estimate of false-positives arising from non-specific hybridization (e.g., in adhesive ‘sticky’ tissue regions).

**False or blank code:** a barcode assigned to neither target nor negative probes, yet decoded for quality control purposes. Detection of these codes provides an estimate of false-positives arising from background fluorescence, optical artifacts, or barcode decoding errors.

**Fluorescent reporter:** a fluorescently labeled oligonucleotide carrying a fluorophore (e.g., organic dyes such as Cy-series or Alexa Fluor dyes) that binds to the encoded readout sequence of a target probe during a given imaging cycle, thereby generating the detectable signal used for transcript identification and localization.

**Fluorescent barcode:** a predefined pattern of positive (*on*) and negative (*off*) fluorescent signals across one or more imaging cycles that uniquely identifies a molecular target. Each bit represents the expected presence or absence of a fluorescent reporter in a given imaging round and channel.

**Fluorescent codebook:** the collection of fluorescent barcodes assigned to measured molecular targets and controls. It represents a subset of the theoretically possible barcode space, with unassigned barcodes serving as false or blank codes for estimating decoding errors and background signal.

**Barcode scalability:** the ability to increase the number of distinguishable molecular targets without proportionally increasing the number of imaging cycles. In conventional cyclic detection, each additional target requires an additional dedicated labeling, limiting the number of measurable targets by the number of available cycles and fluorescent channels. Error-robust barcode designs overcome this limitation by assigning targets to combinations of multiple binary on/off bits across imaging rounds; for example, a 4-color, 16-bit encoding scheme can theoretically generate up to 2^16^ unique binary barcodes, with subsets selected based on Hamming weight and distance to ensure robust decoding.

**Hamming weight** denotes the number of positive, or *on*, bits in a fluorescent barcode, i.e., the number of imaging positions in which any fluorescent reporter is expected to be detected. For example, the barcode 1001 0100 0000 0000 has a Hamming weight of 3. **Hamming distance**, by contrast, counts the bit positions that differ between two barcodes; it quantifies the number of bit edits required to convert one barcode into another. For example, the Hamming distance between 1001 and 1100 equals 2.

**Dense encoding:** an encoding strategy where each target molecule is labeled by one of multiple colored fluorophores in each cycle (e.g., green-yellow-green-red). This design maximizes the number of targets that can be encoded for a given bit budget but is more prone to misidentification and molecular crowding. **Sparse encoding** adopts the opposite strategy, using only a few *on* bits (e.g., 1000 0010 0000 0100 in the same scheme) so that most positions are negative, or *off*. This design reduces optical crowding and improves robustness to false-positive detections, at the cost of encoding fewer targets per bit.

**Field of view (FOV)**: the area of a specimen that a microscope can observe at once in a single image, defined by the lenses and camera sensor size. Multiple FOVs are typically tiled and stitched to capture larger tissue areas while maintaining spatial registration of detected signals. Transcript decoding and cell segmentation may be performed independently within individual FOVs or after stitching across adjacent FOVs, enabling continuous reconstruction and minimizing boundary artifacts.

**Capture area** refers to the total region of a sample-slide for which data can be acquired in a single iST run by tiling multiple FOVs. Increasing the number of FOVs expands spatial coverage but lengthens acquisition time and can lower sample quality. Because the chosen tiling geometry interacts with tissue architecture, coarse coverage can miss rare microenvironments and fine coverage can produce sparse data. Thus, the capture area must balance sample size, imaging throughput, and measurement fidelity to limit spatial sampling bias.

### Box 2: Count modeling

Let *s*_*i*_ ∈ ℝ^2^ be the spatial locations of a set of *i* = 1, …, *n* points (transcripts), assumed to be a marked point process with marks *g*_*i*_ ∈ *{*1, …, *m}* corresponding to gene labels. Let *C*_*k*_ ⊂ ℝ^2^ denote the polygon corresponding to segmented cell *k*, with area 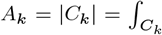, and assume initially that the polygons are pairwise disjoint. For gene *j*, define the aggregated cell-level count

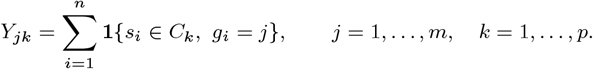

The matrix **Y** = (*Y*_*jk*_) is the gene-by-cell count matrix used in downstream analyses.

**Counts induced by the marked point process**

Represent the marked point process by gene-specific intensity functions *λ*_*j*_ (*s*) with gene *j* ∈ *{*1, …, *m}* so that the intensity measure for gene *j* in a measurable region *B* is Λ_*j*_ (*B*) = ∫_*B*_*λ*_*j*_ (*s*) d*s*. Assuming that the intensity for gene *j* is constant inside cell *k, λ*_*j*_ (*s*) = Λ_*jk*_ with *s* ∈ *C*_*k*_. Conditional on the cell- and gene-specific intensities, the Poisson-process independent-increments property gives

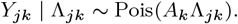

Thus, the point-process model induces a cell-level Poisson model whose offset is the polygon area.

**Gamma-Poisson model across cells**

A convenient model for cell-to-cell heterogeneity is to treat the piecewise-constant intensity as random. For example, following the mixture model introduced by Jones *et al*. [75],

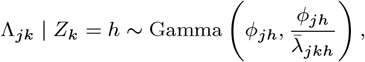

where

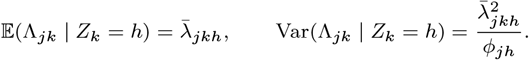

Here, *Z*_*k*_ ∈ *{*1, …, *H}* is a mixture component, potentially representing a cell type, state, or other latent class, with

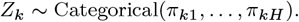

Conditional on *Z*_*k*_ = *h*,

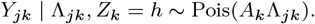

Integrating out Λ_*jk*_ gives a negative-binomial distribution:

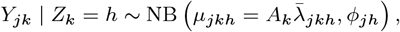

with E(*Y*_*jk*_ | *Z*_*k*_ = *h*) = *µ*_*jkh*_ and 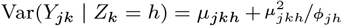. Therefore, this derivation leads to a negative binomial model similar to that of sequencing data, but where the offset relates to the relative size of each cell rather than the total number of sequenced reads.

### Box 3: Expression decomposition

**Decomposition of the mean count**

It is useful to decompose the mean expression in additive terms that correspond to i) biological transcript intensity; ii) technical detection efficiency; and, iii) geometric sampling exposure. Hence, we can write the component-specific expected signal count as

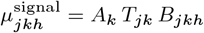

where *A*_*k*_ is the effective observed area or exposure of cell *k*; *T*_*jk*_ is the technical probability or efficiency with which a transcript of gene *j* in cell *k* is detected, decoded, and assigned correctly; and, *B*_*jkh*_ is the underlying biological transcript density for gene *j* in a cell belonging to component *h*.

A flexible decomposition of the **biological component**’s intensity is

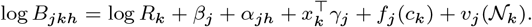

Here, *R*_*k*_ is a cell-level biological scale, such as total RNA content or total transcript density; *β*_*j*_ is the baseline biological abundance of gene *j*; *α*_*jh*_ is the effect of latent cell class *h* on gene *j*; *x*_*k*_ contains observed biological covariates, such as treatment, disease status, genotype, cell-cycle phase, activation, differentiation, or stress; *c*_*k*_ is the cell centroid and *f*_*j*_ (*c*_*k*_) is a spatially varying biological effect; and, *N*_*k*_ describes the neighborhood of cell *k*, and *v*_*j*_ (*N*_*k*_) represents effects of neighboring cell types or the local microenvironment. The scale *R*_*k*_ can depend biologically on cell size, cell volume, ploidy, copy-number variation, cell cycle, transcriptional activity, or metabolic state.

A multiplicative model for the **technical component** in a single tissue and slide is

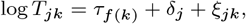

where *τ*_*f* (*k*)_ field-of-view variation; *δ*_*j*_ gene-specific probe hybridization affinity, fluorophore efficiency, and barcode-decoding sensitivity; and, *ξ*_*jk*_ gene- and cell-specific probe effects, such as GC-content and probe-isoform matching. The effective area *A*_*k*_ may differ from the biological cell boundaries. It can include section thickness or depth, clipping by the field-of-view boundary, partial observation of a cell, tissue damage, segmentation erosion or dilation, and inaccessible regions within the mask. Observed area is therefore partly a sampling exposure, while true cell size and volume are biological properties.

**Additive contamination and segmentation mixing**

Not all technical effects are naturally multiplicative. Ambient transcripts, false-positive detections, and cross-cell assignment are additive or mixing effects. A more general observed mean, which allow for contamination due to cell segmentation [75] is

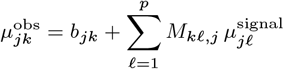

where *b*_*jk*_ is the expected number of ambient or false-positive transcripts assigned to cell *k*; and, *M*_*kl,j*_ is the expected fraction of gene-*j* transcripts originating from cell *l* that are assigned to cell *k*. In an ideal experiment, *M*_*kl,j*_ = **1***{k* = _*l*_*}* and *b*_*jk*_ = 0. Off-diagonal elements of *M* can represent boundary errors, transcript spillover, overlapping masks, incorrect nuclear expansion, optical localization error, and segmentation-induced mixing.

**Identifiability implication**

At the level of the observed count matrix, the principal estimable object is generally the product

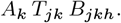

For example, a lower cell-specific detection efficiency *T*_*jk*_ and a lower biological expression level *B*_*jkh*_ have the same effect on the expected count. Similarly, (cell-specific efficiency *×* total RNA content), (gene-specific probe sensitivity *×* baseline gene abundance), and (observed cell fraction *×* true cell size or transcript density) are intrinsically confounded without external information, controls, or normalization constraints. This model is therefore not directly useful in defining a normalization procedure. The decomposition nevertheless gives a complete generative representation of the biological, technical, and geometric mechanisms that can determine the observed gene-by-cell count matrix.

